# The central role of pyruvate metabolism on the epigenetic and molecular maturation of bovine cumulus-oocytes complexes

**DOI:** 10.1101/2022.11.17.516185

**Authors:** João Vitor Alcantara da Silva, Jessica Ispada, Aldcejam Martins da Fonseca Junior, Camila Bruna de Lima, Erika Cristina dos Santos, Marcos Roberto Chiaratti, Ricardo Perecin Nociti, Marcella Pecora Milazzotto

## Abstract

Pyruvate, the end-product of glycolysis in aerobic conditions, is produced by cumulus cells, and is converted in Acetyl-CoA into the mitochondria of both cumulus cells (CCs) and oocytes as a master fuel input for the tricarboxylic acid cycle (TCA). The citrate generated in the TCA cycle can be directed to the cytoplasm and converted back to acetyl-CoA, being driven to lipid synthesis or, still, being used as the substrate for histones acetylation. This work aimed to verify the impact of pyruvate metabolism on the dynamic of lysine 9 histone 3 acetylation (H3K9ac) and RNA transcription in bovine cumulus-oocyte complexes during in vitro maturation (IVM). Bovine oocytes were IVM for 24h in three experimental groups: Control [IVM medium], sodium dichloroacetate [DCA, a stimulator of pyruvate oxidation in acetyl-CoA] or sodium iodoacetate [IA, a glycolysis inhibitor]. Our results show that both treatments change the metabolic profile of oocytes and CCs, stimulating the use of lipids for energy metabolism in the gamete. This leads to changes in the dynamics of H3K9ac during the IVM in both oocytes and CCs with impact on the synthesis of new transcripts in CCs. A total of 148 and 356 differentially expressed genes were identified in DCA and IA oocytes groups, respectively, when compared to the control group. In conclusion, disorders in pyruvate metabolism during maturation stimulate the beta-oxidation pathway, altering the mitochondrial metabolism, with consequences for the mRNA content of bovine oocytes.

## Introduction

Histone acetylation is one of the most important and dynamic epigenetic mechanisms in mammals, contributing to the establishment and maintenance of a permissive environment for transcription (NG HH and Bird, 1999; Tuner, 2002; Gujral, et al., 2020). The reduction of DNA-nucleosome interactions promoted by this epigenetic mark reduces the barriers that prevent transcription factors recruitment (Gujral, et al., 2020). During the oocyte epigenetic maturation, an overall reduction in histone acetylation occurs, carried out by histone deacetylases – HDACs (Pontelo, et al., 2021), leading to the repression of transcription and subsequent chromatin remodeling that precedes the resumption of meiosis (Araujo, et al., 2014).

Changes in the acetylation levels of histone were related to negative effects on mouse oocytes, with an impact on the acquisition of meiotic competence and a consequent decrease in embryonic development (MA, et al., 2013). A similar behavior was observed in human oocytes treated with HDAC inhibitors, leading to an increased incidence of aneuploidies and decreased oocyte quality (Huang, et al., 2012). On the other hand, bovine oocytes matured in the presence of HDAC inhibitors have increased embryonic development rates possibly due to the attenuation of meiotic progression (Saraiva, et al., 2022). Also, changes in acetylation levels collaborate with the functioning of oocyte transcriptional machinery, promoting increased activity of certain groups of genes that may improve the dynamic of epigenetic maturation, such as HAT1 and HDAC1 (Pontelo, et al., 2020).

Besides the epigenetic remodeling, oocytes must undergo a series of alterations known as oocyte molecular maturation, which includes the transcription, storage, and processing of RNAs to support the early embryonic development until the major genome activation, around 8-16 cells in bovine embryos (Graf, et al., 2014). The RNA synthesis occurs mainly during oocyte growth, persisting during oocyte maturation and ending only when chromosomes are fully condensed and become inactive (Sarwar, et al., 2020). During these processes, the cumulus cells (CC) play an essential role as they directly transfer RNAs through transzonal projections and, indirectly, influence the levels of RNA polymerase II transcripts and, consequently, the expression of other genes (Regassa, et al., 2011; Macaulay, et al., 2016). Furthermore, oocytes matured in the absence of CCs present decreased fertility potential, compromising embryo formation, confirming that the CCs exert an essential function for the oocyte during in vitro maturation (IVM) (Uhde, et al., 2018).

In addition to the synthesis and transfer of transcripts, CCs transfer to the oocyte regulatory proteins and metabolites through gap junctions (Sutton, et al., 2003; Turathum, et al., 2021), influencing, for instance, oocyte maturation. Oocyte meiotic resumption requires glycolytic activity (Turathum, et al., 2021, Su, et al., 2009), however, the female gamete itself has a low capacity to metabolize glucose, relying on metabolites transferred by CCs (Sutton, et al., 2003, Turathum, et al., 2021, Wen, et al., 2020). More specifically, glucose is metabolized through the glycolytic pathway in CCs, ending up with pyruvate synthesis and transport to support energy production in oocytes (Andrade and Poehland, 2021). The pyruvate can be oxidized to acetyl-CoA, which subsequently participates in the tricarboxylic acid cycle (TCA) in the mitochondria, contributing to the generation of important substrates for oxidative phosphorylation and, consequently, ATP synthesis (Milazzotto, et al., 2020). The citrate generated in the TCA cycle can also be directed to the cytoplasm and converted back to acetyl-CoA. This acetyl-CoA can be used for lipid synthesis or serve as the substrate for histone-acetyltransferases (HDAC) enzymes to promote the acetylation of nuclear histones (Mcdonell, et al., 2016). In this sense, the metabolic modulation of oocyte maturation may affect histone acetylation both in cumulus cells and oocytes, impacting the molecular control of this complex.

In the present work, we demonstrate how the modulation of pyruvate production/metabolism can induce epigenetic changes in oocyte and cumulus cells. More specifically, the reduction in glycolysis and, consequently, pyruvate production or the increase in pyruvate conversion to acetyl-CoA has consequences on histone acetylation and transcriptional control of bovine oocytes during IVM.

## MATERIAL AND METHODS

### Chemicals and reagents

All chemicals were obtained from Sigma-Aldrich (St. Louis, MO, USA), unless otherwise specified. Tissue culture media (TCM) 199-Hepes and sodium bicarbonate, and fetal calf serum (FCS) were obtained from Gibco (Grand Island, NY, USA).

### Oocyte collection and in vitro maturation (IVM)

Cumulus-oocyte complexes (COCs) were obtained by aspiration of 2-4 mm follicles from bovine slaughterhouse-derived ovaries. COCs were selected and washed three times in collection media (tissue culture medium 199 (TCM-199) with 4-(2-hydroxyethyl)-1-piperazineethanesulphonic acid (HEPES), 10% fetal bovine serum (FBS), 0.2 mM pyruvate and 1.25 mg mL^-1^ gentamicin). Only Grade I and II COCs (Sarwar, et al., 2020) were selected and IVM was performed in groups of 25 COCs in 90 μL droplets of maturation medium (TCM-199 with bicarbonate supplemented with 10% FBS, 0.5 mg mL1 FSH (FolltropinV; Bioniche), 100 IU mL1 human chorionic gonadotropin (hCG; Chorulon; Merck Animal Health) and 1.0 mg mL1 oestradiol) under mineral oil at 38.5°C with 5% CO_2_ in air and high humidity.

### Dose-response of metabolic modulators

For metabolic modulation, oocytes were submitted to IVM in maturation medium supplemented with different doses of dichloroacetate (DCA) or iodoacetate (IA). DCA is an inhibitor of pyruvate dehydrogenase kinase (PDK), a negative regulator of pyruvate dehydrogenase (enzyme from the pyruvate dehydrogenase complex (PDC)). PDC is located at the mitochondrial matrix, acting to convert pyruvate to acetyl-CoA, so the inhibition of PDK increases acetyl-CoA production (ED, et al., 2010). IA acts inactivating glyceraldehyde-3-phosphate dehydrogenase (GAPDH) and, consequently, the conversion of glyceraldehyde-3-phosphate to 1,3-bisphosphoglycerate, diminishing pyruvate synthesis in glycolytic pathway (Xie, et al., 2016).

Bovine COCs were submitted to IVM in medium supplemented with 0.1 mM, 0.5 mM, 1.5 mM and 5 mM of DCA and 0.1 μM, 0.5 μM, 1.5 μM and 5 μM of IA for 24 h. After this period, COCs were stained for mitochondrial membrane potential (MMP) and oocyte maturation. Doses for each modulator were chosen by the following criteria: 1) expected effect on the cumulus cell’s mitochondrial activity and 2) absence of negative effect on nuclear maturation. Results of dose-response are described on *Supplementary Figure 1*.

### Metabolic modulation of IVM

To assess the impact of DCA and IA on IVM, bovine COCs were subjected to IVM in maturation medium (Control group) and in maturation medium supplemented with 2 mM of DCA (DCA group) or 5 μM of IA (IA group). COCs from each group were collected at 4, 8, 16 and 24 h after the beginning of the treatment for analysis. Immature COCs were also collected and submitted to analysis (IM group), as presented in the experimental design (Figure 1). For oocyte evaluation, cumulus cells were mechanically removed by successive pipetting in maturation media and the oocytes were collected and stored until analysis.

**Figure 1.**
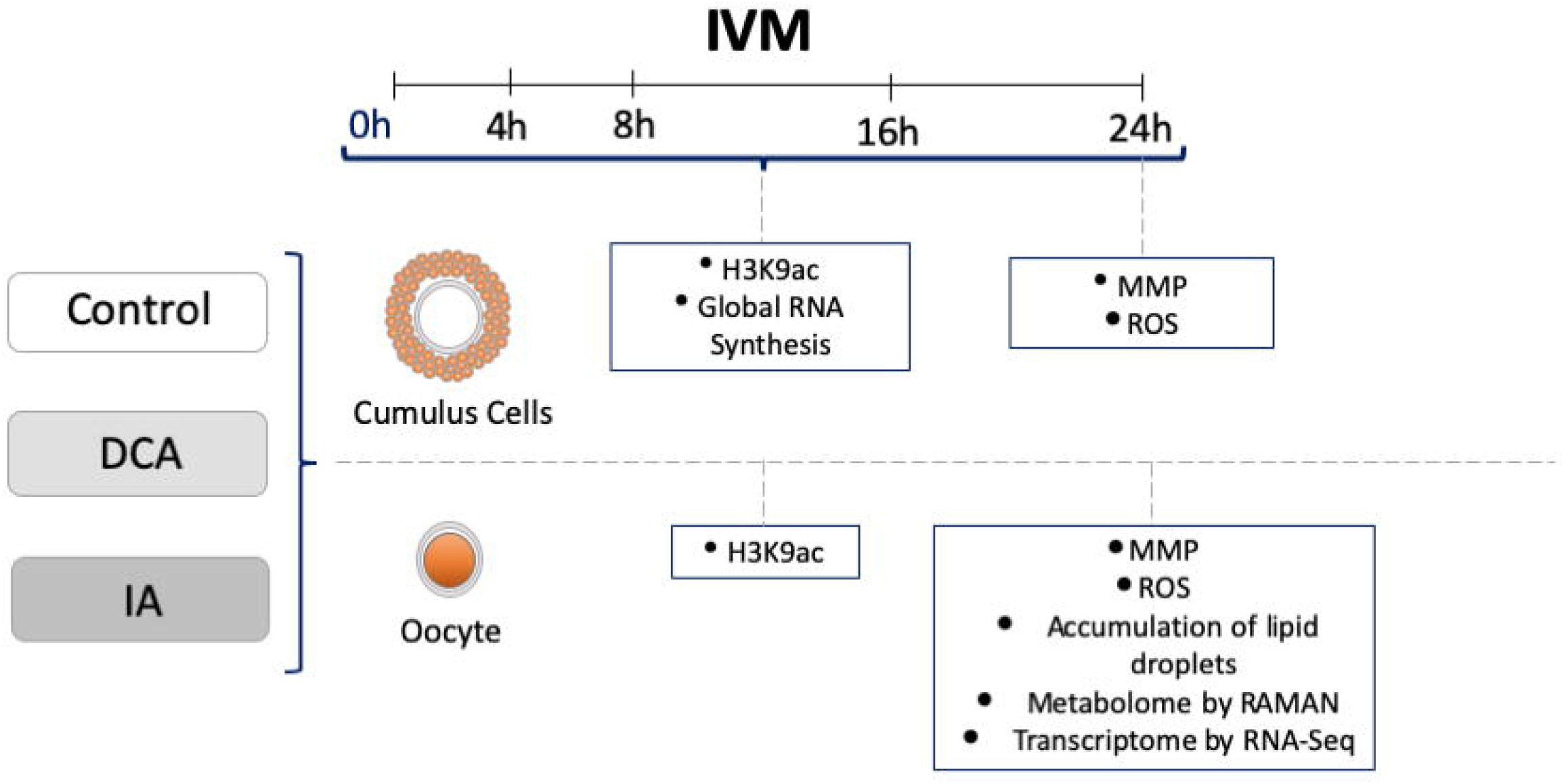
Experimental design. Cumulus-oocyte complexes (COCs) were obtained from bovine slaughterhouse-derived ovaries and submitted to IVM in maturation media supplemented with dichloroacetate (DCA) or iodoacetate (IA). COCs from each group were collected after 4, 8, 16 and 24h and CCs were analyzed by means of H3K9ac and synthesis of new transcripts. After 24h, CCs were assessed for MMP and ROS content. Oocytes from each group were also collected after 4, 8, 16 and 24 h and analyzed by means of H3K9ac. After 24h, oocytes were assessed for MMP and ROS content, metabolome by RAMAN spectroscopy, quantification of lipid droplets and transcriptome by RNASeq.

### Relative quantification of Reactive oxygen species (ROS) and Mitochondrial Membrane Potential (MMP) in cumulus cells

COCs were collected and analyzed after 24 h of IVM by means of ROS content (n=60; minimum of 20 per group) with CellROX^®^ Green Reagent (CRG) (Thermo Fisher) and MMP (n=60; minimum of 20 per group) with MitoTracker^®^ Red CMXRos (Thermo Fisher). COCs from all groups were transferred to a 50 μL drop of PBS containing CRG (5μM) + MitoTracker^®^ Red CMXRos (0.05 μM) and incubated sheltered from the light for 30 minutes in the same conditions of IVM. After incubation, COCs were washed in PBS, mounted on slides in glycerol drops and immediately evaluated under fluorescence microscopy (Leica Microsystems DM16000 B [MitoTracker: Ex/Em 538–617 nm, CRG: Ex/Em 495–519 nm]. Stained COCs were individually photographed at 40x magnification using the Leica Application Suite (LAS) software (Leica Microsystems Brazil).

### Relative quantification of Lipid droplets (LD), Reactive oxygen species (ROS) and Mitochondrial Membrane Potential (MMP) in oocytes

Oocytes were collected and analyzed after 24 h of IVM by means of LD content with Nile Red staining (catalog number: 72485) (n= 45; minimum of 15 per group), ROS content (n=60; minimum of 20 per group) with CellROX^®^ Green Reagent (CRG) (Thermo Fisher) and MMP (n=60; minimum of 20 per group) with MitoTracker^®^ Red CMXRos (Thermo Fisher). After IVM, the oocytes were fixed by 30 minutes in 4% paraformaldehyde solution and stored in the refrigerator (4°C). Subsequently, the denuded oocytes were incubated for 1 hour in 90 μL of IVM medium containing 15 μg/ml Nile Red probe. Then, the samples were washed in 0.1% PBS/PVP. The oocytes were transferred to slides and analyzed under the Leica Microsystems microscope DM16000 B [LD: Ex/Em 552-636 nm]. For MMP and ROS, oocytes from all groups were transferred to a 50 μL drop of PBS containing CRG (5 μM) + MitoTracker^®^ (0.05 μM) and incubated sheltered from the light for 30 minutes in the same conditions of IVM. After incubation, oocytes were washed in PBS, mounted on slides in glycerol drops and immediately evaluated under fluorescence microscopy (Leica Microsystems DM16000 B [MitoTracker: Ex/Em 538–617 nm, CRG: Ex/Em 495–519 nm]. Stained oocytes were individually photographed at 40x magnification using the Leica Application Suite (LAS) software (Leica Microsystems Brazil).

### Metabolomic analysis of oocytes by Raman spectroscopy

Oocytes collected after 24 h of IVM were washed in PBS and immediately analyzed. The Raman spectroscopy setup details and analysis were described elsewhere (Dos Santos, et al., 2016). Briefly, ten Raman spectra were collected per oocyte (n=120; minimum of 10 per group; total of 3 replicates) by using a Triple T64000 Raman Spectrometer (Horiba Jobin-Yvon S.A.S., France) with microanalysis option and CCD detector 1024 × 256—OPEN-3LD/R with quantum response of ~40%. The excitation laser was 532 nm focused on a spot with 5-mW power. The spectra were acquired by using a plan achromatic 50× objective glass (0.20 mm/NA ¼ 0.50) with time of exposure of 20 s, and confocal aperture in 6.5 μm. Data were plotted using the Origin 8.0 software (OriginLab, Northampton, Massachusetts) and pre-processed to remove spikes. The pre-processed data were further analyzed using the Spectrograpy 1.2.15 software to identify the peaks and noise of the spectra with the following parameters: Limit 5% and Prominence 3. Peak attributions were done according to J. De Gelder et al., 2007.

### Immunofluorescence assays for H3K9ac in COCs and oocytes

For acetylation of H3K9 immunofluorescence staining, COCs and oocytes from different groups were collected at 0 (immature oocytes), 4, 8, 16 and 24 h of IVM. Samples were fixed in 4% paraformaldehyde (w/v in PBS) for 30 minutes at RT, permeabilized for 30 minutes with 0.2% TritonX-100 in PBS solution at RT, blocked with blocking solution at 37°C for 3 h and incubated for 2 h with primary antibody (H3K9ac – Abcam ab12179, 1:300) and 1 hour with secondary antibodies (Donkey anti mouse – Abcam ab150109, 1:300) diluted in blocking solution at RT. COCs and oocytes were then stained with bisbenzimide (Hoechst 33342) for nuclear identification, placed in slides and analyzed in a fluorescence microscope [Leica Microsystems DM16000 B coupled with fluorescence filters A4 (blue - excitation/emission 350–490 nm) and L5 (Green - excitation/emission 512-542 nm) and Leica model DFC365 FX camera].

### CCs transcriptional activity detection

The temporal detection of CCs RNA transcription was performed by using Click-iT RNA Alexa Fluor 594 Imaging Kit (ThermoFisher Scientific), according to the manufacturer’s instructions. Briefly, 1 hour before collection of COCs, 1 mM of modified nucleotide ethynyl uridine was added to the medium from different groups. After incubation, COCs were transferred to the *Click-iT reaction cocktail* and incubated for additional 30 minutes. COCs were then fixed in 4% paraformaldehyde and stained with bisbenzimide (Hoechst 33342) for nuclear identification, placed in slides and analyzed in a fluorescence microscope [Leica Microsystems DM16000 B coupled with fluorescence filters A4 (blue - excitation/emission 350–490 nm) and L5 (Green - excitation/emission 485-520 nm) and Leica model DFC365 FX camera].

### Analysis of the global transcriptomic profile of oocytes by RNASeq

cDNA synthesis and amplification were performed using the SMART-Seq HT Kit (Takara Bio Inc) and pools of 5 oocytes (^~^0.1 ng of total RNA) per group per replicate (3 replicates), following the manufacturer’s recommendation. Next, cDNA was purified using AMPure XP Beads (Beckman Coulter), and cDNA concentration and profile were determined using Qubit dsDNA High Sensitivity (ThermoFisher Scientific) and Bioanalyzer High Sensitivity DNA Kit (Agilent), respectively. Libraries were prepared using the Nextera XT DNA Library Prep (Illumina) as recommended by the SMART-Seq HT kit. Libraries were assessed using Qubit and Bioanalyzer, before sequencing on the NextSeq 2000 (Illumina) considering 100 bp pair-end reads.

The quality of the reads was assessed using FastQC (http://www.bioinformatics.babraham.ac.uk/projects/fastqc/). The 100 bp reads were mapped using Star [Dobin, et al., 2013], and identification and quantification were performed using ARS-UCD1.2 (Ensembl and NCBI) as a reference genome using the featureCounts implemented in Rsuberead package [Liao, et al., 2014, Liao, et al., 2019] for gene count. Once the genes were identified, differential expression analysis was performed between groups using DESEq2 [Love, et al., 2014] considering a padj<0.1 and an absolute log2Folchange >0.6. Additionally, we considered genes as differentially expressed if they were exclusive, expressed in one group (expressed in all samples from the same group), and not expressed in the other group (zero counts in all samples from the same group) within comparison and using the function filterByExpr from edgeR package [Robinson, et al., 2010]. We estimate the hub genes using CeTF [Biagi, et al., 2021] based on RIF—Regulatory Impact Factor and PCIT—Partial Correlation and Information Theory [Reverter, et al., 2010; Reverter, et al., 2008]. Gene ontology analyses were performed using clusterProfiler [Yu, et al., 2012] and pathways were explored using Pathview[Luo, et al., 2013]. Data were visualized using R software, in which we primarily observed the classification, intensity, and difference in expression between groups. DEGs were subjected to functional annotation and the enrichment analysis was done by submitting the lists of DEGs to ClusterProfiller. The annotation was conducted with Bos taurus background, and the grouping was performed based on “biological processes’’.

### Statistical analysis

Immunofluorescence images were analyzed by Fiji software. Fluorescence quantification for oocyte MMP, ROS and LD were determined taking into account the number of pixels within the area of each oocyte. For Raman analysis, after peak attribution, the relative intensities were compared. Quantification of H3K9 acetylation in oocytes was determined as the number of pixels within the area of each nucleus. For the quantification of fluorescence in cumulus cells experiments, the oocyte region and approximately 5 layers of the innermost cumulus cells were discarded from the analysis. The total pixel count of this region was quantified and then normalized by COC area.

The mean fluorescence intensity of each image was quantified after area normalization, subtracting the background value. These values and the values of peak intensity for Raman spectroscopy were submitted to an outlier detection test (Grubbs’ test with Alpha = 0.05) and the normality test of D’Agostino & Pearson. In case of normal distribution data, student’s T-Test was performed for comparisons between control and treatments. In case of non-parametric data, the Mann-Whitney test was performed. Nuclear maturation data were analyzed by Chi-square test. All statistical analyzes were performed in a Prism^®^ environment 5.01 (Graphpad Software. San Diego, CA) with a significance level of 5%.

## Results

### The modulation of pyruvate metabolism during IVM alters the histone acetylation profile and the transcriptional dynamic in CCs

The cumulus cells play essential roles during IVM, including the metabolization of glucose through the glycolytic pathway, allowing the transfer of important intermediates for the acquisition of oocyte competence. The modulation of the pyruvate metabolism may also affect the CC metabolism, including the mitochondrial function with consequences to the molecular dynamics of these cells. To verify that, initially we assessed the mitochondrial membrane potential (MMP) and reactive oxygen species (ROS) content in CCs treated with both modulators, DCA and IA. The DCA treatment led to an increase in MMP without interfering with ROS content while CCs incubated with IA had lower MMP and ROS content after 24 h IVM (Figure 2 A-C).

**Figure 2.**
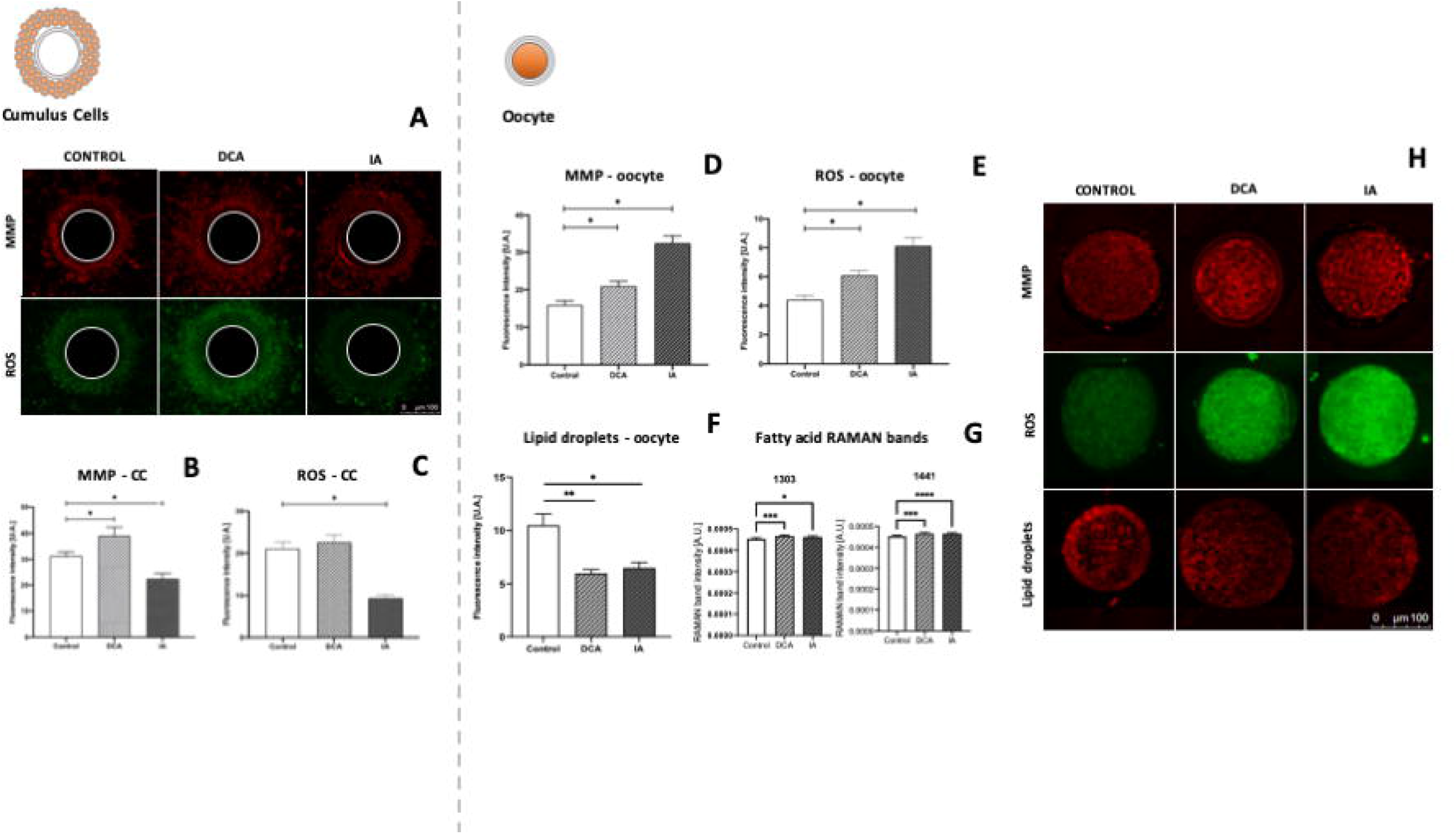
Metabolic analysis of CCs and oocytes. A – representative images of COCs stained for MMP and ROS content analysis. B – Mitochondrial membrane potential of CCs. C – ROS content in CCs. D - Mitochondrial membrane potential of oocytes. E – ROS content in oocytes. F – quantification of lipid droplets of oocytes. G – Fatty acid RAMAN bands intensities of oocytes. H - representative images of oocytes stained for MMP, ROS content and lipid droplets. Data are represented as mean ± S.E.M. *Represents p < 0.05. **Represents p < 0.01. *** Represents p < 0.001

DCA and IA were also capable of altering the H3K9ac profile and the transcriptional dynamics in CCs. DCA increased the levels of H3K9ac in all analyzed time points, and these were followed by an increase in new transcripts synthesis at 8 h and 16 h of IVM (Figures 3B and 3E). Surprisingly, cells treated with IA also had an increase in H3K9ac at 8 h and 16 h (Figures 3A-3C). However, IA led to a decrease in transcript synthesis at 4 h, followed by an increase after 16 h of treatment (Figures 3E and 3F).

**Figure 3.**
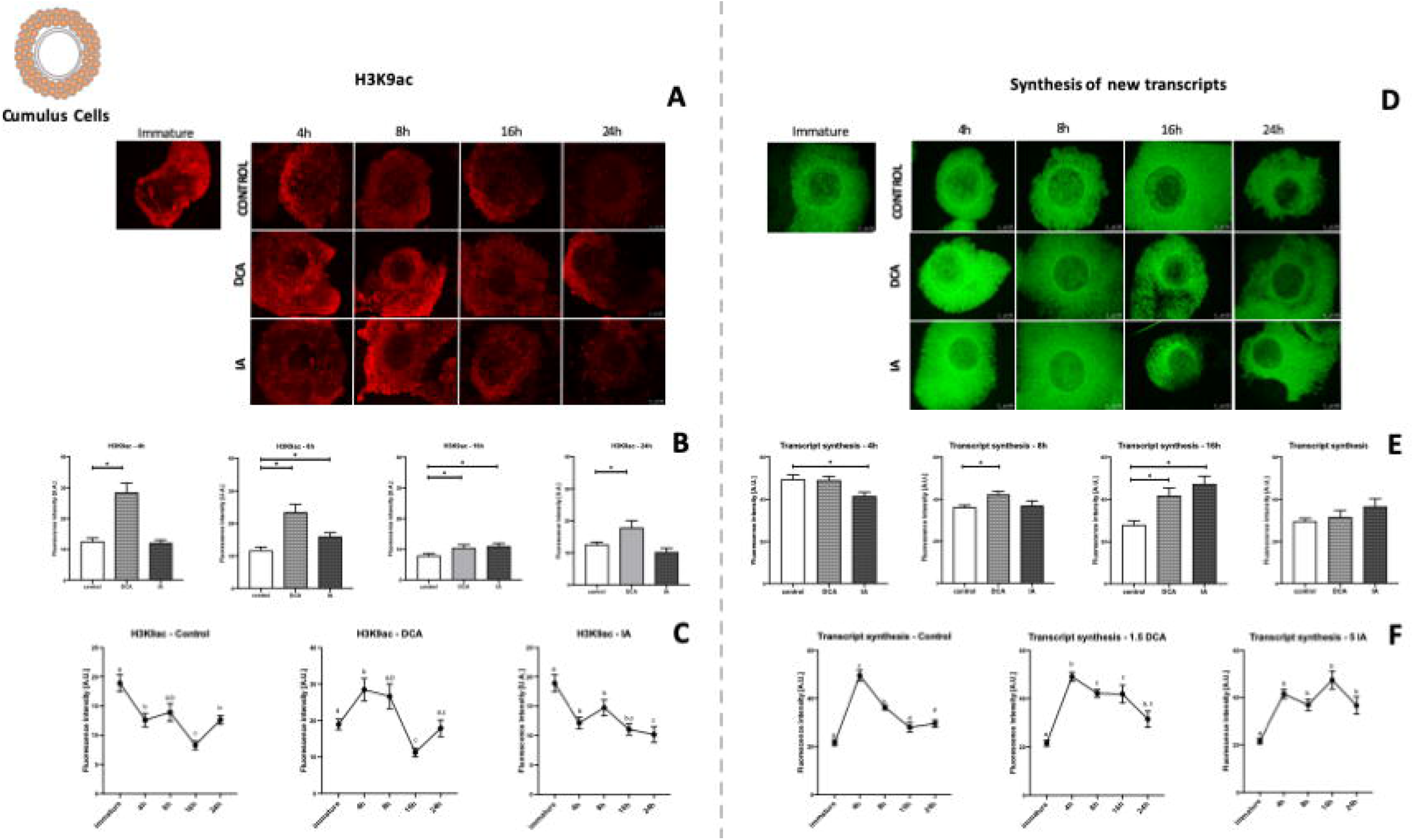
Representative images (original magnification 100×) and Fluorescence intensity levels of H3K9ac (A-C) and synthesis of new transcripts (D-F) in cumulus cells from the control, DCA and IA groups. B – H3K9ac intensity at 4, 8, 16 and 24 h of IVM. D – H3K9ac dynamics during maturation in control, DCA and IA groups. E – Synthesis of new transcripts at 4, 8, 16 and 24 h of IVM. F – Transcriptional dynamics during maturation in control, DCA and IA groups. Data are represented as mean ± S.E.M. *Represents p < 0.05. **Represents p < 0.01. *** Represents p < 0.001 comparing each treatment with the control group. Different letters represent differences regarding each time point for each group.

### Changes in the metabolic parameters of CCs generate consequences on metabolic machinery and oocyte molecular control

The communication established between CCs and oocytes is crucial for the success of oocyte maturation because it is through it that there is the transfer of metabolites, especially pyruvate, which will ensure the functioning of metabolic machinery and the consequent acquisition of oocyte energetic homeostasis (Uhde, et al., 2018). After observing the important metabolic and molecular alterations in CCs, metabolic and molecular parameters in the gamete were assessed (Figure 2D-H).

As in CCs, MMP and ROS content were initially evaluated, and as expected, an increase was observed in the DCA group (Figure 2A-C). Surprisingly, different from what was seen in CCs of this group, there was an increased MMP and ROS content in group IA (Figure 2A-C), suggesting that negative regulation of the glycolytic pathway in CCs could lead to the mobilization of other substrates for the maintenance of energy homeostasis, such as the use of fatty acids through beta-oxidation (Figure 2B). In fact, a decrease in the accumulation of cytoplasmic lipids in the DCA and IA group were observed (Figure 2F). Furthermore, RAMAN spectroscopy analysis revealed greater intensity in bands related to free fatty acids in both groups when compared to the control group (Figure 2G).

Corroborating the hypothesis that metabolic alterations may lead to changes in the epigenetic profile of gametes, both treatments led to changes in H3K9ac reprogramming dynamics that presented lower levels than the control group in the DCA group at 4 and 8 h and in the IA group at 16 h after the onset of IVM (Figure 4A). At the end of IVM, the oocytes are expected to lose the H3K9ac marks, compatible with the resumption of meiosis. Interestingly, after 24 h of IVM part of the oocytes from both groups remained with acetylated H3K9, although at a lower level than that found in immature oocytes (Figure 4A-C).

**Figure 4.**
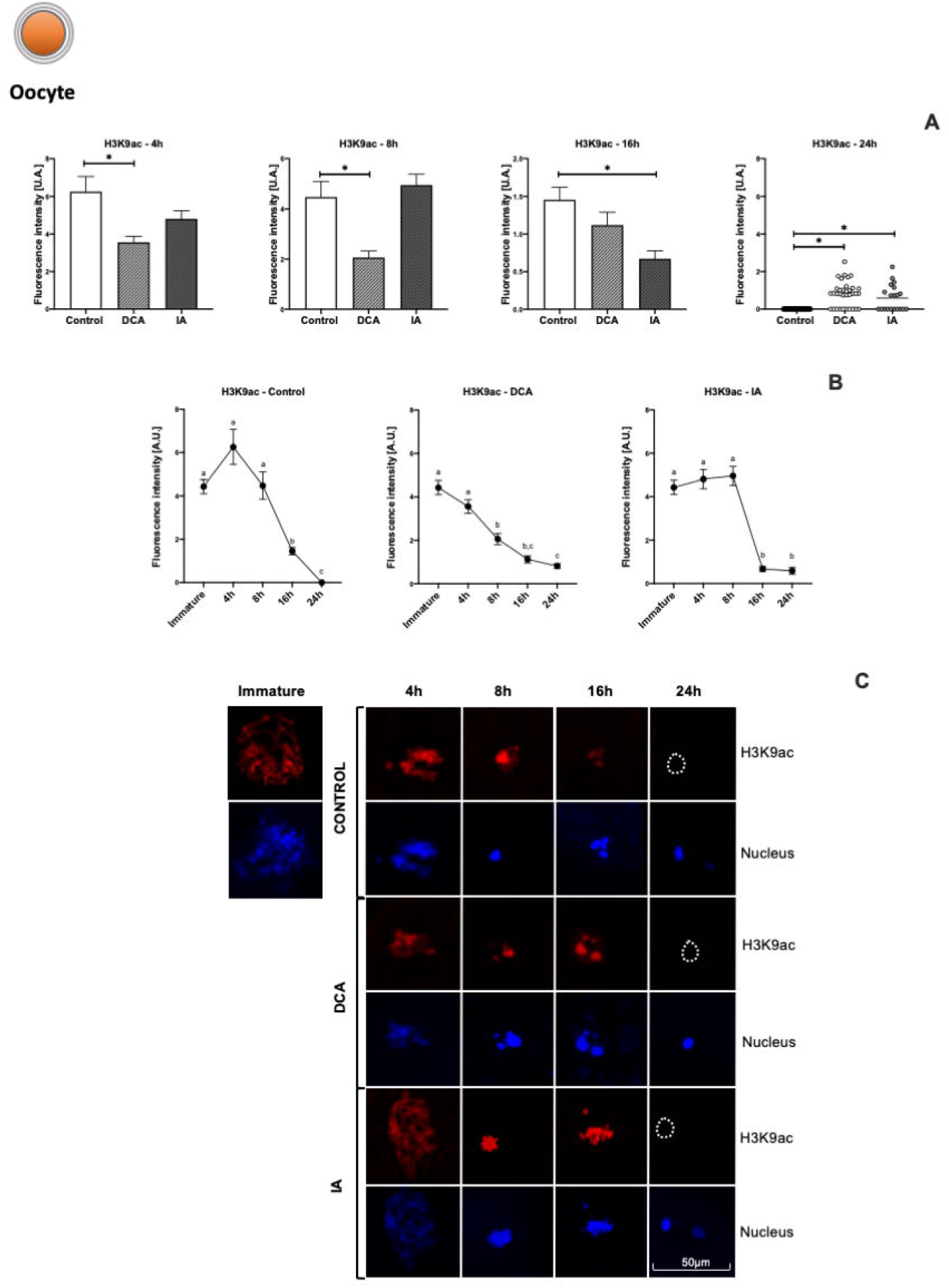
Fluorescence intensity levels and representative images (C - original magnification 50×) of H3K9ac in oocytes from the control, DCA and IA groups. A – H3K9ac intensity at 4, 8, 16 and 24 h of IVM. B – H3K9ac dynamics during maturation in oocytes from control, DCA and IA groups. Data are represented as mean ± S.E.M. *Represents p < 0.05. **Represents p < 0.01. *** Represents p < 0.001 comparing each treatment with the control group. Different letters represent differences regarding each time point for each group.

### Changes in the metabolic and molecular profile of COCs lead to different transcriptional profiles in bovine oocytes

Considering that metabolic alterations in COCs altered the reprogramming dynamics of oocyte H3K9ac, the characterization of the global transcriptomic profile of this cell was performed to investigate the effect of these modifications on the content of mature oocyte transcripts. In the comparison of the control group with the DCA group, 148 differentially expressed genes (DEGs) were observed, 50 genes upregulated and 96 downregulated, while the IA group presented 356 DEGs, of these 152 were upregulated and 203 downregulated (Figure 5).

**Figure 5.**
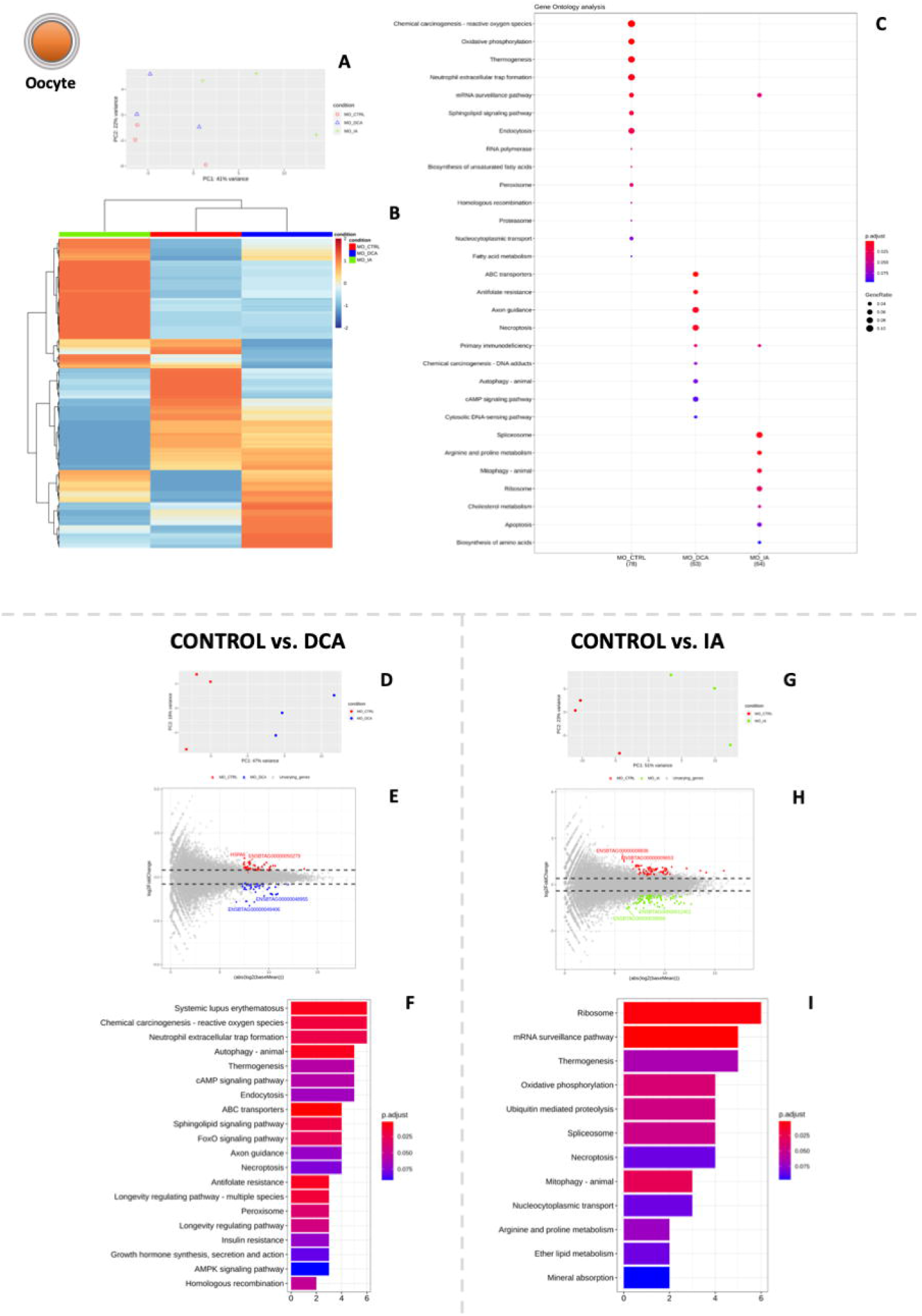
Molecular evidence for DCA, IA and control group. A - PCA analysis of transcripts from control, DCA and IA groups. B – Heat map of DEGs and linkage analysis of groups for the comparison control vs. DCA vs. IA. The most affected biological processes are indicated in (C). D - PCA analysis of transcripts from control vs. DCA group. E – Volcano plot of DEGs for the comparison control vs. DCA. The most affected biological processes are indicated in (F). G - PCA analysis of transcripts from control vs. IA group. H – Volcano plot of DEGs for the comparison control vs. IA. The most affected biological processes are indicated in (I).

The analysis of the biological processes differentially regulated in the DCA vs. control group revealed important pathways for oocyte competence such as “reactive oxygen species”, “autophagy”; AMPK and cAMP signaling”, and “insulin resistance” (Figure 5F and supplementary table 1). The comparison between IA and the control group presented DEGs related to “mRNA surveillance”, “oxidative phosphorylation”, “mitophagy”, and “amino acids metabolism” (Figure 5I). This result evidences the important relationship between epigenetic maturation and its consequent control in oocyte transcriptional machinery and transcript storage.

## Discussion

In this work, we demonstrated that the modulation of pyruvate metabolism generates consequences in the metabolic and transcriptional machinery of CCs, with impacts on the molecular maturation and epigenetics of oocytes during IVM. Pyruvate metabolism is the main source of oocyte energy, however, due to the lack of metabolic mechanisms, the oocyte depends on CCs for glucose metabolization and the consequent transport of pyruvate to the oocyte cytosol (Turathum, et al., 2021; Su, et al., 2009; Andrade, et al., 2021). In the oocyte, pyruvate tends to be metabolized to acetyl-CoA and used in the TCA cycle for the synthesis of ATP and other important metabolites required for the regulation of cellular processes in the gamete.

DCA is a pharmacological modulator that stimulates the activity of the enzyme pyruvate dehydrogenase, increasing the conversion of pyruvate to Acetyl-CoA. Previous studies with mouse embryos have shown that the supplementation of 1 μM of DCA during embryonic culture resulted in higher mitochondrial membrane potential, pyruvate metabolization, and ATP concentration, with a decrease in ROS (Mcpherson, et al., 2014). In addition, Penzias, et al (1993) reported that, unlike IA, DCA has a positive impact on the embryonic development of rodents. In the present work, there was an increase in MMP in COCs. However, regarding ROS amounts, oocytes presented an increase of these molecules while CCs have no significant difference. We believe that this difference in ROS concentrations is due to the fact that oocytes are more susceptible to oxidative stress during IVM (Combelles, et al., 2009) and that CCs have a higher concentration of antioxidants, such as glutathione and melatonin, when compared to oocytes (Cetica, et al., 2001; Mengden, et al., 2020). Finally, the oocytes treated with DCA also seem to use lipid metabolism for energy acquisition, and it is important to emphasize that this process is associated with increased ROS in mouse oocytes and mammalian cells (Ordonez, et al., 2014; Mihalas, et al., 2017).

Recent studies have shown that the metabolism and acetylation of histone are closely connected, with glucose metabolism acting as the main source of substrates for histone acetylation (Wellen, et al., 2009; Yucel, et al., 2019). However, it has recently been shown that lipid metabolism may be the main metabolic pathway related to the maintenance of histone acetylation since fatty acid supplementation in cell culture responds to 90% of the acetyl groups that are used for acetylation (ED, et al., 2010). We demonstrated that the TCA cycle stimulation led to the increase of H3K9ac in all periods analyzed in CCs, reflecting the key role of glucose and lipid metabolism in maintaining this epigenetic mark, with consequent changes in the transcriptional dynamics of these cells. However, in oocytes, we observed the increase of H3K9ac only in some gametes at 24 h post IVM. We believe that this behavior may be associated with the adequate action of HDAC1 enzymes and, mainly, HDAC2 (high expression during oocyte maturation) which acts by deacetylating histones during IVM (MA, et al., 2016).

There are two mechanisms by which histone acetylation regulates gene transcription. The first one is based on the reduction of positive charges in histones promoted by acetylation, which reduces the strong interactions between histone proteins and DNA, exposing the DNA to transcription factors (Bannister and Kouzarides 2011; Gujral, et al., 2020). The second one is by the creation, stabilization, or break of interaction regions between chromatin and regulatory proteins, such as transcription factors or proteins that act on chromatin condensation (Milazzotto, et al., 2020; Santos-Rosa and Caldas 2005). Regardless of the mechanism, there is evidence that exposes this important relationship between histone acetylation and transcriptional cell machinery. Recently it has been reported in human endometrium cells that histone acetylation levels are related to transcriptional cell activity and cellular phase (Munro, et al., 2010).

IA is a pharmacological modulator responsible for inhibiting the glycolytic pathway by the inactivation of glyceraldehyde 3-phosphate dehydrogenase (GAPDH), a glycolytic enzyme. It has been reported that supplementation with both IA and DHEA (another glycolytic pathway inhibitor) during bovine IVM resulted in a decrease in blastocyst rates (Lipinska, et al., 2021). It is also known that the decrease in glycolytic pathway activity affects the quality of oocyte maturation (Melanie, et al., 2012). Our results demonstrated that IA acts by decreasing the MMP of CCs, indicating the importance of these cells in pyruvate metabolism. Surprisingly, we demonstrated an increase in MMP in oocytes. This event may be associated with the fact that CCs create a microenvironment that acts by protecting the female gamete from molecules that can generate negative effects on oocyte maturation, such as IA (Turathum, et al., 2021). However, IVM is characterized as a stage of great energy demand, and it has been reported that oocytes submitted to environmental metabolic stress tend to activate other metabolic pathways to ensure energy homeostasis (Andrade, et al., 2021).

During oocyte maturation, bidirectional communication between oocytes and CCs ensures the transport not only of pyruvate but also of lipids (reference). Lipids are an important metabolic alternative due to their cost-effective ATP production. These molecules are metabolized to fatty acids and transported to the mitochondria using the enzyme carnitine palmitoyltransferase (CPT) so that they can be oxidized by B-oxidation (Andrade, et al., 2021; Dunning, et al., 2010). However, in IVM, the oocyte tends to stock lipids as a form of prevention of lipotoxicity; but when initiated the metabolization of these biomolecules generates increased ROS and consequent cell death (Ordonez, et al., 2014). Here we demonstrated that the inhibition of the glycolytic pathway in COCs led to the decrease of cytoplasmic lipid droplets and the increase of fatty acids in the oocyte, with a consequent increase of ROS in this gamete. This supports the hypothesis of a metabolic alternative use, B-oxidation, for the acquisition of energy homeostasis. The increased use of lipid metabolism using L-carnitine supplementation has also been observed in mouse embryos that interestingly generated the improvement of oocyte quality and the increase in the number of cells of the inner cell mass (Dunning, et al., 2010). Finally, B-oxidation also proved to play an important role in nuclear maturation and the acquisition of competence in bovine oocytes (Paczkowski, et al., 2013).

Acetyl-CoA derived from pyruvate or lipid metabolism can be used in the synthesis of ATP or as substrate by histone acetyltransferase in maintaining the acetylation levels of histone (Pontelo, et al., 2020). Histone acetylation is known as a permissive epigenetic mark of gene transcription. However, for the correct meiotic progression, IVM is characterized by a decrease in the levels of histone acetylation (Ordonez, et al., 2014). Surprisingly, we demonstrated that treatment with IA generated an increase in H3K9ac in some oocytes at 24 h, suggesting that some gametes suffered adequate deacetylation of histone. However, the increase in H3K9ac observed in other oocytes may be associated with decreased oocyte quality due to the formation of chromosomal aberrations that have already been observed in the oocytes of humans treated with deacetylase inhibitors (Pontelo, et al., 2021).

Due to the loss of histone acetylation levels in IVM and the consequent decrease in transcriptional permissibility, oocyte maturation is characterized by a moment of rapid transcription and translation (Luong, et al., 2020). Moreover, for the adequate activity of transcriptional machinery, the oocyte receives transcripts from the CCs through cytoplasmic projections. A study reported that the transcripts transported to the oocyte are related to fundamental processes for oocyte maturation, such as cell cycle regulation, DNA repair, chromatin organization, and energy metabolism (Wyse, et al., 2020). However, there are reports that this connection between oocytes and CCs is locked until 8 h after the beginning of IVM (Uhde, et al., 2018). Knowing this, interestingly we could observe an increase in H3K9ac at 8 h, the supposed moment of cytoplasmic bonds closure in COCs, with a consequent increase in transcription synthesis at 16 h, suggesting that acetylation levels are correlated with transcription levels in CCs. Considering the metabolic stress microenvironment, this increase in H3K9ac and transcription may be associated with the attempt of CCs to transcribe molecules that ensure adequate oocyte maturation, because studies have shown that there is a transcriptional dependence between the oocyte and CCs that are determinant for the success of oocyte maturation (Biase and Kimble, 2018).

There is already evidence in cellular models that support the association of energy metabolism and the levels of histone acetylation with the activity of transcriptional machinery (Mcdonnel, et al., 2016; Gao, et al., 2021). These reports corroborate the findings in the present work since the metabolic and epigenetic alterations promoted here by the different treatments are also associated with different transcriptional profiles. The DCA presented transcripts upregulated related to B-oxidation, ACBD7, and genes associated with the maintenance of histone proteins, such as HIST1H3C, HIST1H2BL, and HIST3H2A. There is evidence that transcripts that act in the maintenance of the nucleosome are related to the progression of tumors, such as the findings of HIST1H3C and HIST3H2A (Zao, et al., 2021; Paramashivam, et al., 2015). IA showed downregulated genes related to glucose metabolisms, such as GID4 and SLCA4. Interestingly, SLCA4 has been reported to collaborate with tumor progression by regulating pH in an environment with low oxygen levels (Carroll, et al., 2022). On the other hand, IA induced upregulation of genes associated with lipid metabolisms, such as APOA2, MTTP, CLPS, and TECR. Also, there was a downregulation in transcripts related to embryonic development, such as the PLG that inactivated presents decreased embryonic survival in mice (Ploplis, et al., 1995), and DBX1, which is fundamental for embryonic and fetal development in vertebrates (Karaz, et al., 2016).

Taken together, our results indicate that changes in pyruvate metabolism induce the activation of metabolic pathways for the acquisition of energy homeostasis during maturation. However, this event causes changes in the dynamics of histone acetylation in CCs, with a consequent change in the transcriptomic profile of the oocyte, demonstrating the important relationship between the metabolic and transcriptional machinery of COCs. This work reinforces the importance of studies in metaboloepigenetics to better comprehend the gametes and embryo development and also as a tool for the improvement of the bovine IVP system.

