## Supplementary Table 2 for "The central role of pyruvate metabolism on the epigenetic and molecular maturation of bovine cumulus-oocytes complexes"

### Supplementary table 2 – Differentially expressed genes (DEGs) of CO vs IA comparison.

| GeneID | LogFoldChange | PValue |
|---|---|---|
| ARL6IP4 | 2.470057868201 | 0.0493291468072162 |
| PPCS | 1.0125297070539 | 0.0023832806824189 |
| MOV10 | 1.0325029072932 | 0.00703570481896919 |
| DRAM2 | 1.0540280667981 | 0.0186959392568578 |
| CD164L2 | 1.0599235409248 | 0.0133766741465196 |
| LOC104973208 | 1.1384413027489 | 0.0249588605078733 |
| ENSBTAG00000023464 | 1.2079114742634 | 0.00403334756841737 |
| LOC112442278 | 1.2559446548829 | 0.00284296362819712 |
| ENSBTAG00000004925 | 1.4292025246355 | 0.0446704064229655 |
| ENSBTAG00000008836 | 1.5214995751357 | 0.000298190581023792 |
| ZNF770 | 1.5627858043701 | 0.0231397467328437 |
| SLC25A43 | 1.7899943027601 | 0.0189083092108252 |
| ENSBTAG00000055013 | 1.7982847385443 | 0.00600707532558889 |
| COL12A1 | 3.5585398330315 | 0.040958737071969 |
| ZBTB26 | 4.0706483146311 | 0.0136219961612608 |
| MIR2887-2 | 1.00330778986231 | 0.0120282714348134 |
| ENSBTAG00000011654 | 1.00359002665774 | 0.00170452373892715 |
| LOC107133207 | 1.00562998557163 | 0.012828714950532 |
| LOC112445012 | 1.00649223923648 | 0.0193151217065487 |
| LOC112442207 | 1.00683686838058 | 0.00333381705188188 |
| PTCD1 | 1.01511222817058 | 0.00873319560214087 |
| PPIL3 | 1.01628775702375 | 0.00397143905315128 |
| ENSBTAG00000051088 | 1.02002360607156 | 0.000246198347164889 |
| GUK1 | 1.02279237491005 | 0.0480083570278997 |
| LOC616720 | 1.02866716415153 | 0.00285154360594084 |
| TMTC3 | 1.02951393178869 | 0.000495689250783545 |
| ENSBTAG00000011671 | 1.03263113905247 | 0.0157795321013649 |
| OLAH | 1.03542997746265 | 0.00214659204939071 |
| H2AFX | 1.03989820956724 | 0.0149474817592361 |
| LOC100140550 | 1.04068920424638 | 0.00223715044400465 |
| SLC6A4 | 1.04110218652044 | 0.010245652848053 |
| ENSBTAG00000051889 | 1.04153853831295 | 0.000713772994305415 |
| LOC101907545 | 1.04521399633937 | 0.0324547270580982 |
| ENSBTAG00000054176 | 1.04689341979635 | 0.00114353745016762 |
| CDC37 | 1.05104970306976 | 0.0432654508362876 |
| MCOLN2 | 1.05438319183939 | 0.013054609486499 |
| ENSBTAG00000050174 | 1.06816032701892 | 0.00413350774245176 |
| HSD17B11 | 1.07217272105274 | 0.00592482824945133 |
| LOC112447479 | 1.07517610159663 | 0.000679493029055227 |
| METTL4 | 1.07610597360384 | 0.0385894942808922 |
| MAP3K1 | 1.08063526702358 | 0.010619713441218 |

| | | |
|---|---|---|
| ARMCX5 | 1.08218524722445 | 0.00596650638483217 |
| HIST1H2AJ | 1.08573699229878 | 0.00179539224269238 |
| LOC104970926 | 1.08747225066758 | 0.00171859535970364 |
| ENSBTAG00000046373 | 1.08801021753611 | 0.00177530589927074 |
| ZNF214 | 1.08951068293881 | 0.00319921685479485 |
| CENPX | 1.09061601237062 | 0.018288924059765 |
| UQCRC1 | 1.09066157129069 | 0.0189220028317208 |
| GTPBP10 | 1.09163660063957 | 0.00158060654128678 |
| SWSAP1 | 1.09214153336071 | 0.0496028926919128 |
| CEP19 | 1.09381054685217 | 0.0306628162777079 |
| GSTP1 | 1.09652491301217 | 0.0244086684783202 |
| TLR2 | 1.10365706772164 | 0.00779102497325679 |
| ENSBTAG00000030305 | 1.10749119632071 | 0.000187887321098913 |
| HPGD | 1.11174074606491 | 0.0433416347333625 |
| LOC524176 | 1.12397080819439 | 0.0141190619549894 |
| LIN37 | 1.13011548535149 | 0.00852077439667795 |
| LOC100848921 | 1.13307489260652 | 0.00262026392743935 |
| USE1 | 1.13578554835274 | 0.00890785618573125 |
| CKB | 1.13699404144461 | 0.0224229061894921 |
| ENSBTAG00000053833 | 1.14618489085078 | 0.0233258552108451 |
| ENSBTAG00000049323 | 1.15136221926294 | 0.0102534179433769 |
| ACADL | 1.15299954460296 | 0.0170077825187424 |
| LOC101904701 | 1.16958725148566 | 0.000121941790232867 |
| C5H12orf60 | 1.17809101925668 | 0.0013817257394504 |
| PGBD5 | 1.18713188838824 | 0.0201256055921842 |
| UBE2U | 1.19751608745865 | 0.000722671985828893 |
| ICOS | 1.20307431437069 | 0.00691791324810214 |
| TSC22D1 | 1.20542874829028 | 0.00731809700750689 |
| ATP5F1D | 1.20990595673725 | 0.0304079403320021 |
| ENSBTAG00000048837 | 1.22109072438952 | 0.00721895022581818 |
| LOC112442340 | 1.22289187010915 | 0.00418890385730299 |
| ENSBTAG00000045985 | 1.22716278791808 | 0.000499722347981162 |
| RFESD | 1.22927798734895 | 0.0228582272366513 |
| MDGA1 | 1.23280233649863 | 0.00782736969726207 |
| ENSBTAG00000033360 | 1.23297685540366 | 0.0165437925909297 |
| SLCO1C1 | 1.23975907220407 | 0.00188541190285366 |
| ENSBTAG00000044025 | 1.25202878724753 | 0.0400251334710275 |
| TSPAN6 | 1.25510428338506 | 0.000326688288088329 |
| LACTB2 | 1.25697043119437 | 0.000138076203047188 |
| ENSBTAG00000022278 | 1.30046474523398 | 0.0125769001052384 |
| EFHC2 | 1.33577766888753 | 0.0346450431587401 |
| CXHXorf38 | 1.34045420382471 | 0.0451001831032769 |
| LOC618696 | 1.35858150844161 | 0.0396049029666975 |
| TMEM52 | 1.35987887527989 | 0.00829947164523942 |

| | | |
|---|---|---|
| ENSBTAG00000043482 | 1.36781691245887 | 0.0175273899396633 |
| ENSBTAG00000050863 | 1.37712465059983 | 0.0350522553317347 |
| ENSBTAG00000055267 | 1.37989323344019 | 0.0112330411806431 |
| MS4A13 | 1.39188377848234 | 0.0257495724507743 |
| CAPS2 | 1.39263308050471 | 0.0439678554175342 |
| H4 | 1.39458773446012 | 0.00201854807631727 |
| TECR | 1.40584566830312 | 0.0149441856728777 |
| LOC777593 | 1.41681805235688 | 0.00214165818465296 |
| HIST1H3C | 1.41720603739518 | 0.0363976450909525 |
| NMRAL1 | 1.51262846176608 | 0.00305122533816298 |
| KLRG2 | 1.54287171425196 | 0.0458280516594899 |
| EYA4 | 1.57638506748661 | 0.0411458141257946 |
| ENSBTAG00000038422 | 1.57729132707605 | 0.000971128541598323 |
| EFCAB13 | 1.58842033518805 | 0.0189406897844192 |
| LOC101903261 | 1.61089086373595 | 0.0294862779207165 |
| LOC789715 | 1.61964582003424 | 0.037727779029762 |
| SYT4 | 1.63431164006705 | 0.0251698131631935 |
| LOC101902831 | 1.68912633523791 | 0.00456447959996275 |
| ANKRD35 | 1.71100945898099 | 0.0285862067496045 |
| TMEM71 | 1.73775719264303 | 0.0462854217173535 |
| NAA11 | 1.74375690959534 | 0.0326716043851104 |
| SLC18B1 | 1.74595642703836 | 0.0463694155706876 |
| LOC112449049 | 1.74934376209946 | 0.0219295649524726 |
| PAFAH1B3 | 1.75317393006759 | 0.0214431316318941 |
| ZNF711 | 1.76576080615619 | 0.0177998766685703 |
| LOC112445964 | 1.79230736847732 | 0.005433405967421 |
| OVOL3 | 1.79416678603999 | 0.0266051791449562 |
| UST | 1.85799651692566 | 0.00139141700101703 |
| LOC112449115 | 1.90510591794626 | 0.0212954258165472 |
| LOC107133022 | 1.90652034722388 | 0.0368250317356762 |
| ENSBTAG00000054456 | 1.97287077066907 | 0.0290054205322789 |
| LOC112446641 | 1.98442660357925 | 0.0410987300820344 |
| DTYMK | 2.05559890080504 | 0.040220948483451 |
| ISG15 | 2.17179614233058 | 0.030300189071457 |
| DAGLB | 2.17485366295354 | 0.0111535839552658 |
| KIF19 | 2.20867188543033 | 0.0309370876870009 |
| LOC107132623 | 2.31879084390541 | 0.0193913886589578 |
| GABRA1 | 2.54141118471133 | 0.0266096308570395 |
| LOC104972276 | 2.57935491209376 | 0.0184810858027402 |
| LOC112449112 | 2.59617667961663 | 0.00816740386851865 |
| MEIOB | 2.62237604416474 | 0.0258350831387494 |
| LOC100848940 | 2.62366651341121 | 0.00466648201931314 |
| MTTP | 2.69339465531119 | 0.00252166822463316 |
| FATE1 | 2.71179483716218 | 0.0358132345590826 |

| | | |
|---|---|---|
| MPV17L | 2.80465926469654 | 0.0327211618805593 |
| LOC617189 | 3.12442186998899 | 0.0368605938621135 |
| LOC781298 | 3.13744915121727 | 0.0326171592201751 |
| LHFPL5 | 3.20578791207887 | 0.00732995569675479 |
| LOC112448164 | 3.31177452427669 | 0.0293425840394743 |
| DDX49 | 3.53315458630224 | 0.00205455933798757 |
| HRH3 | 3.55374480923927 | 0.0433621951159586 |
| CD80 | 3.55632210645346 | 0.0437919509746548 |
| ENSBTAG00000013305 | 3.61290956375271 | 0.0341620938349204 |
| LOC511401 | 3.64481612476525 | 0.0337189022469799 |
| ITGA10 | 3.75321275357298 | 0.0252052613300476 |
| ENSBTAG00000046633 | 3.76692461082414 | 0.0410113056343695 |
| APOA2 | 3.96491118613269 | 0.0469702755638601 |
| KLK6 | 3.98643721348346 | 0.0480514577932866 |
| ENSBTAG00000051842 | 3.98643721348346 | 0.0480514577932866 |
| DERL3 | 4.02072450263102 | 0.0448709277696118 |
| LOC112449281 | 4.18426263376341 | 0.0178985593733481 |
| CLPS | 4.32422721702938 | 0.0196153690812313 |
| FCGR2A | 4.41893514416404 | 0.0171998493892659 |
| SNCB | 4.47523985746721 | 0.0131957396672771 |
| APOH | 4.59361573619867 | 0.00919306950784142 |
| IFI44 | 5.03433256400293 | 0.00254150820688749 |
| IFI44L | 5.65003730727755 | 0.000280716570282068 |
| ENSBTAG00000050384 | -4.98740668549259 | 0.00194099092759986 |
| SPSB1 | -4.58654793316618 | 0.00217131229774498 |
| SDHAF1 | -4.32964658633361 | 0.0179893186404389 |
| LOC112447932 | -4.21150298915789 | 0.0267623763116699 |
| RUSC2 | -4.19829195552422 | 0.0291766452657059 |
| SERPINH1 | -4.19829195552422 | 0.0291766452657059 |
| TYR | -4.07254511297069 | 0.0361931692917693 |
| LOC104970180 | -3.91557361931347 | 0.0494226483255801 |
| PLEK | -3.91151166559818 | 0.0466052221062206 |
| ENSBTAG00000051430 | -3.90832367727693 | 0.0469013047707465 |
| C18H19orf84 | -3.90034208989847 | 0.0476463779428499 |
| IL6 | -3.78660333822504 | 0.0225416982775623 |
| ENSBTAG00000025814 | -3.77543057226613 | 0.0220382513913321 |
| TMEM202 | -3.77543057226613 | 0.0220382513913321 |
| ENSBTAG00000050198 | -3.68944451601976 | 0.0393567255506659 |
| ENSBTAG00000052996 | -3.65703868635086 | 0.00197317665402579 |
| LOC101902430 | -3.65703868635086 | 0.00197317665402579 |
| LOC112445863 | -3.60394724764561 | 0.00992434826795863 |
| GSTK1 | -3.59619225137714 | 0.0375396541522213 |
| CTXN2 | -3.51751359102798 | 0.00925512551436984 |
| SEL1L2 | -3.47215038706903 | 0.0473005940543588 |

| | | |
|---|---|---|
| UBE2L6 | -3.47053532719201 | 0.0412370870074201 |
| BMX | -3.40224724413775 | 0.0229791953670303 |
| ENSBTAG00000054985 | -3.36419123757542 | 0.0233618502274585 |
| LOC112445162 | -3.06034376395808 | 0.0106444713641847 |
| ENSBTAG00000039357 | -3.01446044163387 | 0.000180623295217822 |
| ENSBTAG00000054273 | -2.93831588331873 | 0.013691897695606 |
| TMEM176A | -2.79969740749494 | 0.0108807267127003 |
| ENSBTAG00000023318 | -2.77527709376484 | 0.0131830103789905 |
| CSTA | -2.71693188354711 | 0.0328911398737715 |
| ENSBTAG00000052249 | -2.71063370308658 | 0.0160780627989153 |
| PALMD | -2.64789557474883 | 0.0196329641350519 |
| IL27RA | -2.64556209009013 | 0.0215662810426183 |
| LOC112443242 | -2.63473764559736 | 0.00651931277446588 |
| LOC104972629 | -2.61151969596463 | 0.00254139703968381 |
| SVOP | -2.58833015552174 | 0.00545239932122868 |
| ENSBTAG00000046974 | -2.58023088864485 | 0.0466455405750349 |
| GLI3 | -2.52215398022493 | 0.0297912195077952 |
| KCNB2 | -2.45904796269843 | 0.00997412375396252 |
| ENSBTAG00000009620 | -2.44033791386103 | 0.0149011674451256 |
| RADX | -2.43361661927302 | 0.0287018444949093 |
| CXHXorf57 | -2.43361661927302 | 0.0287018444949093 |
| ENSBTAG00000031764 | -2.42235549297984 | 0.0102565841126033 |
| ENSBTAG00000008020 | -2.35431811765769 | 0.0443063597922728 |
| LOC104971508 | -2.30820298117761 | 0.00321717142179485 |
| LOC509283 | -2.30572531453006 | 0.0263875573062265 |
| WDR27 | -2.23152891803455 | 0.0118114610403647 |
| ENSBTAG00000051928 | -2.21808856007682 | 0.00128266264554042 |
| THBS2 | -2.18346938686417 | 0.00124544596716961 |
| TACR1 | -2.15198989112082 | 0.0079637854168367 |
| CDHR1 | -2.12729417793274 | 0.00952263305147634 |
| NUP210 | -2.09789403944642 | 0.0219271954466297 |
| LOC112445945 | -2.09684561084508 | 0.0023040159431965 |
| ENSBTAG00000054825 | -2.09048863241196 | 0.0124638364113537 |
| NDRG1 | -2.07372165877529 | 0.0312218808732642 |
| ADH4 | -2.05554840071594 | 0.00757146417853878 |
| LOC107133071 | -2.04149687359533 | 0.0315438823908815 |
| RMDN2 | -2.02015386819784 | 0.0437901595954836 |
| SCN4B | -1.98602286729894 | 0.00958325528036138 |
| PLAU | -1.97356663397364 | 0.01894429642545 |
| ENSBTAG00000049284 | -1.97121054422954 | 0.000906932405555578 |
| LENG8 | -1.95654219429821 | 0.00399781884026545 |
| ENSBTAG00000052977 | -1.92492537287947 | 0.00110391698856916 |
| NKX3-1 | -1.91102065222851 | 0.0036617307500676 |
| ENSBTAG00000053024 | -1.87461817354808 | 0.000395958136954219 |

| | | |
|---|---|---|
| ENSBTAG00000052792 | -1.78875169998549 | 0.00124996040886413 |
| ENSBTAG00000051171 | -1.75070324855408 | 0.0253580109400293 |
| CAMK1D | -1.74267726237306 | 0.00412593006669971 |
| LOC112448933 | -1.73562191787088 | 0.00846742707655418 |
| ENSBTAG00000051106 | -1.73226647203015 | 0.0103639072562297 |
| FAM199X | -1.71861921221889 | 0.0379199706480002 |
| LOC615959 | -1.70140822000315 | 0.000257220995641304 |
| LOC112444750 | -1.70034902837821 | 0.00507355478000523 |
| ENSBTAG00000045876 | -1.67669544873012 | 0.0253739701876876 |
| ENSBTAG00000047836 | -1.64915655126529 | 0.0476939570152844 |
| LOC107132435 | -1.64720464979472 | 0.00953381087192031 |
| CD3D | -1.59210100613197 | 0.0257697550716637 |
| LOC112449395 | -1.58418374015923 | 0.00089049173183765 |
| PRR12 | -1.57379497407283 | 0.00209210170554749 |
| LOC104974297 | -1.56335805645892 | 0.0295569914589031 |
| VSTM1 | -1.55020073865585 | 0.0353444856215127 |
| ENSBTAG00000014454 | -1.52384058734037 | 0.0136993375121118 |
| TBC1D25 | -1.52265287723876 | 0.00244951079779304 |
| PLG | -1.52258595716171 | 0.0153611859874822 |
| LOC112445796 | -1.50848450497528 | 0.000274985512640851 |
| SPAG8 | -1.50430877438176 | 0.0274257674140075 |
| ATP6V1B1 | -1.49016865566861 | 0.0153614391887403 |
| ENSBTAG00000026011 | -1.48989851423378 | 0.000136529905760276 |
| ENSBTAG00000053958 | -1.48218845690174 | 0.00798570076479996 |
| ATF6B | -1.47903500755938 | 0.000855218540751452 |
| PAM | -1.47587363957882 | 0.0143918754773387 |
| RFX1 | -1.47081416641259 | 0.0393991772003662 |
| LOC112448316 | -1.46737646178251 | 0.0368093413004131 |
| FAM83F | -1.46682826838598 | 0.0268003111570979 |
| ENSBTAG00000038238 | -1.46304392990841 | 0.0492431703013291 |
| COL16A1 | -1.46036044228097 | 0.0471546531131305 |
| COMMD7 | -1.45967558180597 | 0.0168957900081831 |
| ENSBTAG00000049575 | -1.45474912457617 | 0.00299735439923703 |
| ENSBTAG00000033169 | -1.44867258814813 | 0.0454653590750114 |
| STOX2 | -1.44846582169693 | 0.0125595437959969 |
| ENSBTAG00000051478 | -1.43837240544656 | 0.0249377458875637 |
| LOC104976269 | -1.43125461410168 | 0.00324615465690472 |
| LOC101903877 | -1.42489418208277 | 0.0018398778017682 |
| ZCCHC12 | -1.42044769402125 | 0.00582334324131164 |
| ENSBTAG00000031483 | -1.41552940087156 | 0.000418129605761867 |
| ENSBTAG00000011161 | -1.39854056820783 | 0.0119835669457712 |
| TGFB2 | -1.39264214316366 | 0.0350571740736995 |
| ZFHX2 | -1.36223653814736 | 0.0303429676338054 |
| JUND | -1.35533149926357 | 0.00191934186152569 |

| | | |
|---|---|---|
| ENSBTAG00000007354 | -1.35285178645701 | 0.00315446515275109 |
| LOC531196 | -1.34459489735054 | 0.00473324177822944 |
| GSK3A | -1.33973972921889 | 0.0026354655698775 |
| ENSBTAG00000050598 | -1.33067643692799 | 0.00493140897403043 |
| ENSBTAG00000005866 | -1.30421123874015 | 0.0157651018001161 |
| ZNF81 | -1.30421123874015 | 0.0157651018001161 |
| SLC2A4 | -1.29132916382073 | 0.00107929536796711 |
| ENSBTAG00000048688 | -1.28093884614115 | 0.0178451882050837 |
| COL26A1 | -1.27912889205912 | 0.0157274872250613 |
| GID4 | -1.26558618764585 | 0.00318337188226048 |
| ENSBTAG00000047194 | -1.26031636375954 | 0.0267410783000916 |
| C5H12orf75 | -1.25975593840577 | 0.0109457765477708 |
| KLHL3 | -1.25787604785594 | 0.00020684203535679 |
| ENSBTAG00000050594 | -1.25006480425217 | 0.000474666617467175 |
| BCL3 | -1.23851999703859 | 0.0247293506151776 |
| ENSBTAG00000051058 | -1.23451086449177 | 0.00347802866325258 |
| ENSBTAG00000050321 | -1.22857222963045 | 0.016121981792648 |
| ENSBTAG00000032281 | -1.22811073003673 | 0.000208602810676951 |
| ENSBTAG00000049275 | -1.22240191602066 | 0.00650123805876119 |
| LOC100297540 | -1.22114799123834 | 0.00316484076289283 |
| USP29 | -1.22061441754567 | 0.00477727088415568 |
| TSHZ1 | -1.21463818717611 | 0.021125141701232 |
| EGR1 | -1.21187977086438 | 0.00459566497933031 |
| THTPA | -1.20697318139053 | 0.0120231066393635 |
| ENSBTAG00000036858 | -1.20526110003728 | 0.0111235115696102 |
| ENSBTAG00000054695 | -1.20369990787327 | 0.0106784510827894 |
| ENSBTAG00000050450 | -1.20319640430838 | 0.00139624041292627 |
| CRABP2 | -1.19754819843011 | 0.0304041695729994 |
| LOC104973390 | -1.19634392158965 | 0.00512474118791349 |
| ENSBTAG00000020723 | -1.19012557152299 | 0.0185313084129971 |
| DNAH14 | -1.18935915529794 | 0.00498123437968742 |
| ENSBTAG00000047699 | -1.18427521642576 | 0.000379148619888571 |
| PIM1 | -1.18208155121631 | 0.00227882556284865 |
| CACNB4 | -1.17810545639143 | 0.000463819498822713 |
| ENSBTAG00000039218 | -1.17549337165281 | 0.00724579212414231 |
| IL7R | -1.17314672380817 | 0.0156755265330784 |
| LOC104968488 | -1.17288285455648 | 0.0107987290359986 |
| MGAT1 | -1.15598135792971 | 0.0242357533486725 |
| ENSBTAG00000007199 | -1.15440064364172 | 0.000701547191447338 |
| LOC112442867 | -1.15149130797071 | 0.00503756121502207 |
| ADGRG6 | -1.14639575348999 | 0.0427183595867808 |
| TEDC1 | -1.14491988084029 | 0.0102523945831997 |
| LOC100300115 | -1.14217286877286 | 0.0285429006241638 |
| ENSBTAG00000048955 | -1.14209800319993 | 0.00579056654194951 |

| | | |
|---|---|---|
| LOC112446383 | -1.14129677607666 | 0.0251570208686638 |
| ENSBTAG00000031464 | -1.14122847721011 | 0.0193533418151539 |
| HECW2 | -1.13655357687871 | 0.0205772998697093 |
| LOC112449293 | -1.12266988064008 | 0.00992995872787867 |
| ATXN2L | -1.12090837390393 | 0.00326960937982636 |
| NLGN1 | -1.12046445993509 | 0.000522918832027671 |
| ENSBTAG00000049167 | -1.11032280534287 | 0.00459203606134751 |
| TMC5 | -1.10734781287663 | 0.00190767694651791 |
| DBX1 | -1.10597618956355 | 0.000210618039280829 |
| ENSBTAG00000053487 | -1.09935820133514 | 0.0242163726003198 |
| CALN1 | -1.09879804041103 | 0.00377164248175887 |
| TNFRSF21 | -1.09582193644018 | 0.0297142046663847 |
| CDC42BPB | -1.09442480573839 | 0.000575787161680729 |
| ENSBTAG00000050097 | -1.09240401757483 | 0.00243440674763819 |
| ENSBTAG00000053258 | -1.08150051070079 | 0.00879695739867683 |
| LOC509155 | -1.07903698658005 | 0.00246757478524454 |
| LOC101902555 | -1.06902843092115 | 0.00811589559805257 |
| LOC107132714 | -1.06811706207671 | 0.00270524263816689 |
| LOC112445164 | -1.06698759955842 | 0.00153987238525931 |
| LOC781112 | -1.06001243233401 | 0.00887085113280114 |
| KANK2 | -1.05943347350748 | 0.033496562822111 |
| ENSBTAG00000054555 | -1.05572483632492 | 0.0473177963954336 |
| LOC112441471 | -1.04130349082387 | 0.0331701813805086 |
| ENSBTAG00000053048 | -1.03806096060354 | 0.00172968274161525 |
| ENSBTAG00000002900 | -1.03563999446109 | 0.00132118409663465 |
| LOC112449259 | -1.03521395540567 | 0.046713640139975 |
| ENSBTAG00000046004 | -1.03364408565762 | 0.0044795446366497 |
| LOC112444152 | -1.02860202783999 | 0.00183398740202458 |
| DPYD | -1.02829777078818 | 0.0244325064975396 |
| ANKRD1 | -1.02704098014148 | 0.0454891902118786 |
| PCDHA13 | -1.02670693128969 | 0.00394182903500357 |
| ENSBTAG00000051246 | -1.02588786916129 | 0.0422899472769363 |
| ENSBTAG00000037434 | -1.02435891115093 | 0.00858692756333467 |
| GARNL3 | -1.01739287323852 | 0.0424882999721769 |
| ASB16 | -1.01223039040058 | 0.0268185017427748 |
| YBX2 | -1.00853649049107 | 0.00119634413622075 |
| ENSBTAG00000006252 | -1.00715977261655 | 0.00236645856176939 |
| LOC112443475 | -1.00507834950937 | 0.0249023914364598 |
| MSI1 | -1.00125420565133 | 0.0126602220428785 |
| LOC112448609 | -4.3632352133987 | 0.00395552709893797 |
| ENSBTAG00000051456 | -2.3415190298593 | 0.000505353259025423 |
| ENSBTAG00000051063 | -1.5655185434645 | 0.00368937496687592 |
| PAK4 | -1.3665887440578 | 0.0305437927917693 |
| LOC100335990 | -1.2713362847689 | 0.00465104500613581 |

| | | |
|---|---|---|
| LOC101902951 | -1.2371810208056 | 0.00357469527085709 |
| LOC101904674 | -1.2075128076221 | 0.00309561082071691 |
| LOC515601 | -1.1145918768249 | 0.00805771059747571 |
| TFE3 | -1.1128081003024 | 0.000900586869432532 |
| ENSBTAG00000050973 | -1.747630812833 | 0.0067670549843861 |
| ENSBTAG00000008557 | -1.16575155629 | 0.00506489010748845 |
