## Supplementary Table 1 for "The central role of pyruvate metabolism on the epigenetic and molecular maturation of bovine cumulus-oocytes complexes"

**Supplementary table 1 – Differentially expressed genes (DEGs) of CO vs DCA comparison.**

| GeneID | LogFoldChange | PValue |
|---|---|---|
| ACBD7 | 1.03871048329247 | 0.0108281681711515 |
| ENSBTAG00000051113 | 1.04020854262504 | 0.0184345872613525 |
| HIST1H3C | 1.04547409588668 | 0.0364660172185324 |
| SOGA3 | 1.0525154535631 | 0.0404526406933195 |
| HSPA6 | 1.0766231318032 | 0.000616976230116506 |
| ZBTB34 | 1.08146525814988 | 0.0258408454996482 |
| ZBTB18 | 1.10668865781902 | 0.0211957012952721 |
| CA8 | 1.11487413509784 | 0.0111469551516905 |
| ASPN | 1.11534908163239 | 0.0195198332447687 |
| PKHD1L1 | 1.11542008447541 | 0.0410771718092637 |
| NRP2 | 1.16229689982015 | 0.0408398457707883 |
| ACADL | 1.16963778370819 | 0.0141080145765361 |
| LOC527388 | 1.18685694683679 | 0.0216003790163372 |
| ASB4 | 1.22739962198527 | 0.0058611753936615 |
| EFHC2 | 1.25878646535893 | 0.0499045871777638 |
| FARP2 | 1.27034781955624 | 0.0413603932167728 |
| ENSBTAG00000049949 | 1.33366400716744 | 0.0122012068325761 |
| FER1L6 | 1.38950385492748 | 0.0341764634400672 |
| HIST1H3C | 1.41889989504183 | 0.0333799349516626 |
| ENSBTAG00000053101 | 1.42392590172307 | 0.030942877951001 |
| LOC104975042 | 1.42520827678171 | 0.0285833026871617 |
| BBOF1 | 1.45620515438406 | 0.0244478296107939 |
| RAB33B | 1.47251778239729 | 0.00176691501067735 |
| LOC101903018 | 1.50676842196612 | 0.000840025732063247 |
| LOC101903567 | 1.5099529567712 | 0.0348363612287756 |
| PARP6 | 1.53357042736321 | 0.0335929240385878 |
| LOC786372 | 1.57324598631205 | 0.0208193539576569 |
| HIST1H2BL | 1.59022957951205 | 0.00295612769333428 |
| ANKRD34B | 1.62427662591905 | 0.0392365869247503 |
| LOC614095 | 1.66518387002676 | 0.0162103612778817 |
| ENSBTAG00000054433 | 1.68514012184076 | 0.000727304448241783 |
| IL1RN | 1.71494123162328 | 0.0217466859992569 |
| LOC112443013 | 1.88092987231034 | 0.0122385560051757 |
| LOC112443614 | 1.8890708681727 | 0.0354830454680135 |
| LOC104972276 | 2.42021608109131 | 0.0232475670574515 |
| GH1 | 2.50641821032328 | 0.0112399836843755 |
| CTSA | 2.5211674642185 | 0.0255988656178522 |
| RAET1L | 2.6197379569017 | 0.0256718051710997 |
| SGK1 | 2.77959063549838 | 0.00902895749845163 |
| SFXN2 | 2.90247850221125 | 0.0481814734591474 |
| LOC101903068 | 3.02476815927588 | 0.0341754804893405 |

| | | |
|---|---|---|
| HIST3H2A | 3.06931781139279 | 0.00655671304105588 |
| FGD2 | 3.37762685208599 | 0.0122077763298416 |
| HRH2 | 3.55718216295326 | 0.0333595219959977 |
| ENSBTAG00000051070 | 3.56225113922821 | 0.0364087281225081 |
| MC2R | 3.58851845806707 | 0.0357261488910495 |
| ENSBTAG00000032544 | 3.60999033282944 | 0.0301144750137465 |
| ARID3C | 3.92430077829303 | 0.0464794034957148 |
| MIR2299 | 3.98627310833646 | 0.0109154213963196 |
| LHFPL6 | 4.20309708979659 | 0.0235257400166865 |
| PDE11A | -1.00194833641118 | 0.0277706290223565 |
| GRK3 | -1.01614914695188 | 0.0492037007917179 |
| LOC112443751 | -1.02165640144152 | 0.0208032639020132 |
| LOC107132781 | -1.02977695214488 | 0.00874928225756041 |
| ZC2HC1B | -1.04269401899328 | 0.0415314843148953 |
| LOC112445796 | -1.07176320106527 | 0.000312791285736811 |
| TMEM100 | -1.07390634497167 | 0.00729149469230001 |
| FAM19A3 | -1.0874558276622 | 0.0318571036027233 |
| ENSBTAG00000051504 | -1.10452170671572 | 0.0358105975515452 |
| DPYD | -1.10545942923055 | 0.0123363858738736 |
| ANKRD1 | -1.11466150270594 | 0.035104335887566 |
| LOC112449060 | -1.1164461796747 | 0.0414659945072987 |
| XYLT2 | -1.11987203872142 | 0.0499667335893989 |
| CSTB | -1.1287083556531 | 0.0159500863815953 |
| MT1A | -1.14026902457695 | 0.00809780829487614 |
| C24H18orf63 | -1.14132055822489 | 0.0152290193522523 |
| LOC112442733 | -1.17864160475297 | 0.0424446272655627 |
| C5H12orf75 | -1.19306259788056 | 0.0013391756811399 |
| FBXL21 | -1.2000405349246 | 0.021635331589534 |
| LOC112447391 | -1.2057751749891 | 0.0200919520910402 |
| PRR12 | -1.22947069263666 | 0.0214510320479623 |
| CLDN11 | -1.23387706347243 | 0.0254326379144845 |
| LOC521568 | -1.2372509444012 | 0.000498802092246175 |
| POSTN | -1.23769736690971 | 0.00420217363154882 |
| PAM | -1.25590111652498 | 0.0205337693137014 |
| ENSBTAG00000050973 | -1.2610861385829 | 0.0314893959120358 |
| LOC528748 | -1.29789561765018 | 0.0158694291812577 |
| LRFN5 | -1.33028377540127 | 0.0435241757719553 |
| IQCA1L | -1.35184289328624 | 0.000522997052229877 |
| ENSBTAG00000014454 | -1.36723302715932 | 0.0309004513212091 |
| PLS3 | -1.38046031605069 | 0.00539060905973282 |
| LOC781112 | -1.38309159136359 | 0.000175027522075335 |
| LOC112445525 | -1.40924721525625 | 0.00269816375781971 |
| ENSBTAG00000048955 | -1.42022569191108 | 0.000109041396160359 |
| LOC526294 | -1.43968331328501 | 0.0150763216506327 |

| | | |
|---|---|---|
| MUC13 | -1.43996910607345 | 0.0286687261503358 |
| LOC101906052 | -1.46984343482569 | 0.000229876225690267 |
| NTS | -1.49660288936268 | 0.0151272731400704 |
| CNTN4 | -1.5118481057998 | 0.00276129421540499 |
| HSD11B1 | -1.51312307036021 | 0.0476889621429139 |
| LOC104969823 | -1.51595232287652 | 0.0346504782918339 |
| MT1E | -1.51898895259584 | 0.00392399815172301 |
| OTOP1 | -1.56763087861052 | 0.0187194154253255 |
| HEBP2 | -1.61215887701463 | 0.0317981096695596 |
| ENSBTAG00000049406 | -1.62670327434382 | 0.0423738483758824 |
| KRT10 | -1.6361887883608 | 0.00278686012882607 |
| ENSBTAG00000050598 | -1.65334904124252 | 0.0443980066440248 |
| LOC104971296 | -1.69813522614392 | 0.0131579453923612 |
| SCN4B | -1.70408938984334 | 0.0334014014423109 |
| MEFV | -1.72780373296552 | 0.00680464669087502 |
| RHOG | -1.74038746832449 | 0.04249504214115 |
| LOC101910047 | -1.75065610902259 | 0.0391162642137383 |
| LOC101909196 | -1.75559190359819 | 0.0199863885118122 |
| ENSBTAG00000053577 | -1.79918541032056 | 0.00316071980816875 |
| NDP | -1.81287566611843 | 0.0470982904362254 |
| IFI27 | -1.82843835800248 | 0.00405802627928322 |
| SH3BGRL | -1.8523806998412 | 0.0396585241723417 |
| ENSBTAG00000049246 | -1.88470410413063 | 0.0248531689896349 |
| LOC112445532 | -1.93871852990639 | 0.0133663404941602 |
| RGS7BP | -1.97832154144338 | 0.00187115044925275 |
| LRRC27 | -2.00718916566648 | 0.0291008852030758 |
| LOC112442665 | -2.08402655034582 | 0.00900722207914053 |
| LOC100847365 | -2.08701118803294 | 0.0373051170137267 |
| PPP1R27 | -2.09468582450793 | 0.0466873162326851 |
| SVOP | -2.12167868291925 | 0.0332275563507561 |
| LOC112449620 | -2.14787935845298 | 0.0421015517255033 |
| ADH4 | -2.22718006767114 | 0.00232930545027309 |
| LOC112444172 | -2.28618942252065 | 0.0473350126867106 |
| MFSD4A | -2.33303095747796 | 0.022799389970446 |
| LOC107131455 | -2.41850680888506 | 0.0404061026274145 |
| LOC616149 | -2.46556621664673 | 0.00818219970451944 |
| ENSBTAG00000054779 | -2.49934810195676 | 0.0286419377735489 |
| LOC112443242 | -2.50003686959576 | 0.0069805895897554 |
| BEX2 | -2.53956100732598 | 0.0492142099060384 |
| GLI3 | -2.55139248855211 | 0.0259837191373043 |
| LOC112445038 | -2.57843199175651 | 0.00261219846218646 |
| ACSM4 | -2.69130574468756 | 0.0365677711972308 |
| RBM11 | -2.77959244927405 | 0.0269330886034411 |
| CSTA | -2.81148287319877 | 0.0163995569618145 |

| | | |
|---|---|---|
| DCN | -2.82006376985714 | 0.00879574114010167 |
| ENSBTAG00000048363 | -2.96015166520925 | 0.0405841649263948 |
| C12H13orf46 | -3.19197133218437 | 0.023867419898956 |
| ENSBTAG00000052996 | -3.32368134287209 | 0.00691356174108548 |
| AKAP4 | -3.32976398291429 | 0.0087720878685465 |
| LOC787786 | -3.39013761765821 | 0.00336162255809749 |
| LOC101902430 | -3.46380196878378 | 0.00351386027207975 |
| CTXN2 | -3.47030333124351 | 0.0114858560700863 |
| SPSB1 | -3.7786390273032 | 0.0200502842567074 |
| TMEM202 | -3.84412600921601 | 0.0189538994090739 |
| TYR | -3.90657364530016 | 0.0457455556259196 |
| PLEK | -3.98720037713402 | 0.042956286190835 |
| LOC112443342 | -4.03654007652438 | 0.0370318379485097 |
| ENSBTAG00000025814 | -4.11216548507942 | 0.00887983278047812 |
| KCNK6 | -4.1264022309236 | 0.0306041555620157 |
| SDHAF1 | -4.38449104023107 | 0.0165093185912636 |
| LOC783210 | -4.75337186457292 | 0.00474452483329564 |
