## Supplementary figures and images for "The central role of pyruvate metabolism on the epigenetic and molecular maturation of bovine cumulus-oocytes complexes"

### Supplementary figure 1.tiff

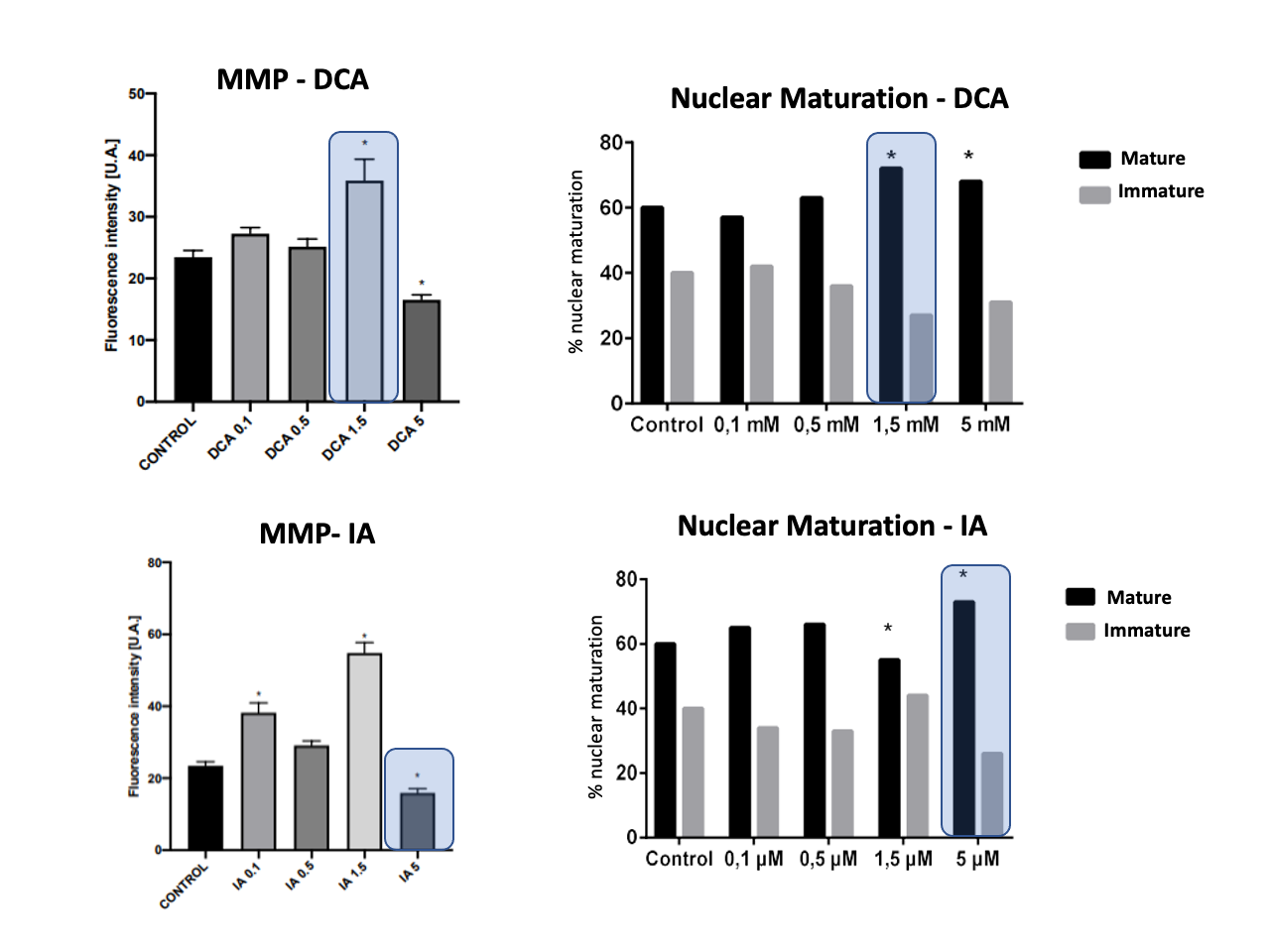
